# Bioremediation of Atrazine and its Metabolite Using Multiple Enzymes Delivered by a *Bacillus thuringiensis* Spore Display System

**DOI:** 10.1101/2023.10.31.565005

**Authors:** Shu-Yu Hsu, Hsin-Yeh Hsieh, George C. Stewart, Chung-Ho Lin

**Affiliations:** Center for Agroforestry, University of Missouri, U.S.A.; School of Natural Resources, University of Missouri, U.S.A.; Bond Life Sciences Center, University of Missouri, U.S.A.; Department of Veterinary Pathobiology, University of Missouri, U.S.A.

**Keywords:** *Bacillus thuringiensis*, atrazine, bioremediation, spore display, multi-enzyme system

## Abstract

Atrazine (ATR), a widely used herbicide in the United States, has contaminated ground and surface water. The persistence in the environment, the carcinogenic characteristic, and the endocrine disruption activity of ATR raises public health concerns. To provide a clean source of drinking water, it is imperative to remediate ATR and its metabolites without toxic by-products. A green technology, *Bacillus thuringiensis* spore-based display system, has been developed to express a high density of enzymes on the spore surface. The *B. thuringiensis* spore can serve as a bioparticle to deliver enzymes to different environments and enhance the stability and shelf-life of enzymes expressed on their surface. A cluster of enzymes, AtzA, AtzB, AtzC, AtzD, AtzE, and AtzF, found in *Pseudomonas sp.* strain ADP, remediates ATR to CO_2_ and NH_3_ in the degradation pathway. AtzA-bearing spores were generated, and they successfully demonstrated the capability of converting ATR to its metabolite, hydroxyatrazine (HA) [Hsieh *et al.* 2020]. In this study, AtzB, a hydroxyatrazine *N*-ethylaminohydrolase that hydrolyzes HA to *N*-isopropylammelide (NiPA), was expressed on the *Bacillus* spores by fusing *atzB* to the gene encoding the attachment domain of the BclA spore surface nap layer protein. The results showed that 1 mg AtzB-bearing spores degraded 50% of HA to NiPA in water within 12 hours. The K_m_ and V_max_ of AtzB-bearing spores with HA as a substrate were 25.28 µM and 1.26 nmole NiPA h^-1^ mg^-1^ spores, respectively. The optimal ratio of 1:2 for AtzA-bearing to AtzB-bearing spores was applied and successfully converted 90% ATR to NiPA in the water after 96 hours of incubation. The surface water with the addition of 34.5 nM (7.5 µg L^-1^) of ATR was treated with a combination of AtzA-bearing and AtzB-bearing spores in a 96-hour time course study. Over 80% of applied ATR in surface water was degraded and ATR concentration was below the United States Environmental Protection Agency maximum contaminant level, 13.8 nM (3 µg L^-1^) within 24 hours. At the end of 96 hours. ATR in surface water was completely converted to NiPA. We have successfully demonstrated the application of multienzyme bioremediation of ATR using the *Bacillus* spore-based display system in the surface water in the laboratory.

## IMPORTANCE

Atrazine contamination in drinking water sources can have adverse effects on human health. Atrazine concentration in surface water, including rivers and lakes, in the United States often exceeded the United States Environmental Protection Agency maximum contaminant level, 13.8 nM (3 µg L^-1^). Novel bioremediation technology is urgently needed to degrade atrazine in drinking water sources. The *Bacillus* spore-based display system is an excellent model to deliver multiple enzymes that are required to completely break down atrazine and its metabolites due to its flexibility in displaying various enzymes, resistance to harsh environments, and enhancement of enzymatic activity. We demonstrated that multiple enzymes delivered by the *Bacillus* spore-based display system completely degraded atrazine in drinking water sources into further non-toxic metabolites. With the successful demonstration of the system in this study, the *Bacillus* spore-based display system could be a solution to degrade other classes of persistent organic pollutants, such as dioxins, in the future.

## INTRODUCTION

Atrazine (ATR), which was banned in the European Union (EU) in 2004, widely contaminated the surface and ground waters in the agricultural production regions in the EU (1, 2), the U.S.(3), Canada (4, 5), and other countries in Asia (6, 7) due to its persistence (half-life > 6 months) and high mobility in the environment (8). Atrazine has been shown to disrupt human endocrine systems, increase the risk of miscarriage and breast cancer, reduce male fertility in humans, and cause intersex in amphibians (9–16). The ATR is one of the world’s most heavily applied herbicides, with up to 36,000 tons applied annually in the U.S. alone since 1990 (10, 17). Greater than 46 nM (10 µg L^-1^, 10 ppb) of ATR has regularly been detected in rivers, groundwater, and drinking water in the U.S. (18), whereas the United States Environmental Protection Agency has set the maximum contaminant level (MCL) at 13.8 nM (3 µg L^-1^, 3 ppb) in drinking water (10, 19). Bioremediation, the use of microorganisms or microbial processes to degrade environmental contaminants, has been recognized as one of the most cost-effective cleanup strategies to remove ATR regarding its efficiency and eco-friendliness (20–27).

The *Bacillus thuringiensis* spores are suitable to deliver enzymes in the environments for bioremediation attributed to the discovery of an enzyme fusion point on the BclA spore surface protein (28)*. Bacillus thuringiensis*, a Gram-positive, rod-shaped bacterium, is commonly found in soil. They sporulate under limited nutrient conditions and the endospores remain dormant in harsh environments. The outermost *B. thuringiensis* spores layer is the exosporium and is composed primarily of proteins and carbohydrates (29). The surface component of exosporium is the hair-like nap that directly contacts the environment (30). The collagen-like glycoprotein, *<u>B</u>acillus* <u>c</u>ollagen-like protein <u>A</u> (BclA), is the major component of the hair-like nap (31, 32). It acts both as a permeability barrier and confers hydrophobic characteristics on the spore (33). The targeting domain within the N-terminal domain of the BclA protein which is critical for the incorporation of the BclA onto the exosporium surface was identified (28). The targeting terminal domain of the BclA can serve as a fusion point to display foreign proteins/enzymes on the spore surface for remediation (34).

The key element to demonstrate the function of the *B. thuringiensis* spore display system is a suitable enzyme for bioremediation of ATR. *Pseudomonas sp*. Strain ADP that was capable of metabolizing >1,000 mg L^-1^ atrazine was first isolated from a herbicide spill (Little Falls, MN, USA) in the 1990s (35–37) when ATR was used as the sole nitrogen source (36). *Pseudomonas sp.* strain ADP carries a plasmid, *p*ADP-1, containing 6 genes, *atzA*, *atzB*, *atzC*, *atzD*, *atzE*, and *atzF*, for ATR degradation. The encoded enzymes sequentially catalyze the degradation of ATR and its metabolites, resulting in the complete mineralization of ATR and its metabolites into CO_2_ and NH_3_ (38). AtzA, the first enzyme in the atrazine degradation pathway (39), has been successfully expressed on the *B. thuringiensis* spore display system (34). The AtzA-bearing spores successfully remediated greater than 90% atrazine in the soil and water from a benchtop study and they have shown higher resistance to the leaching process in the soil than the recombinant AtzA enzyme (34).

With regard to ATR bioremediation in the environment, the *B. thuringiensis* spore-based display system offers several benefits over other conventional bioremediation techniques, such as biostimulation and bioaugmentation. Biostimulation involves the modification of the environment to stimulate the existing bacteria capable of bioremediation. Supporting the existing microbial activity often requires the addition of various forms of limiting nutrients and electron acceptors (40, 41), which is expensive and could promote the growth of other microorganisms that are not ATR-degraders (42). Bioaugmentation, a process that introduces ATR-degrading microorganisms to an ATR-contaminated environment, has been widely studied (43–45). However, the efficiency of the bioaugmentation strategy highly depends upon the ATR-degraders’ ability to compete with indigenous microorganisms, predators, and various abiotic factors (42). The *B. thuringiensis* spore-based display system could minimize the drawbacks of biostimulation and bioaugmentation because no addition of nutrients and maintenance is needed to sustain the enzymatic activity of the expressed enzymes. Potentially, it could also bypass the down-regulation of expression of ATR-degrading enzymes by other nitrogen sources other than atrazine present in the environment (46). Additionally, the *B. thuringiensis* spore-based display system can be engineered to express a variety of genes encoding various enzymes, so it can immediately designed and utilized for multienzyme reactions— the use of several enzymes in reaction steps or cascade reactions without the isolation of intermediates (47)—to remediate ATR. The multienzyme reaction has experienced rapid growth in scientific and industrial applications, such as the synthesis of polymers (48), carbohydrates (49, 50), and plastics depolymerization for plastics remediation (51). The cell-free multienzymes can mediate complicated chemical reactions in a one-pot system and overcome challenges of utilizing microorganisms in the environment, such as their viability, complexity, and physiology (52). The use of multienzymes in a one-pot system, as compared to microbial activity, provides higher product yield, greater engineering flexibility, higher product titer, and faster reaction rate (53).

Remediation of ATR to the further metabolites using the multienzyme reaction has never been explored in the past. Although several studies have utilized AtzA to detoxify ATR to hydroxy atrazine (HA), HA in drinking water could lead to kidney toxicity due to its low solubility in water, resulting in crystal formation and subsequent inflammation (8). Hydroxyatrazine *N*-ethylaminohydrolase (AtzB), the second enzyme in the ATR degradation pathway, was identified as the only enzyme that could transform HA to *N*-isopropylammelide (NiPA)(54). The NiPA can be further transformed into cyanuric acid, which is the central intermediate in ATR metabolism by many bacteria (55). Therefore, AtzB catalyzes a crucial step in ATR remediation. In this study, AtzB-bearing spores were created and characterized. Moreover, the combination of AtzA-bearing and AtzB-bearing spores and their substrate/metabolite dynamics were examined. Finally, to validate the efficacy of the developed novel system in treating the contaminated drinking water sources, the multiple enzymes delivered by the *B. thuringiensis* spore display system were applied to treat the atrazine-contaminated surface water. With the successful demonstration of the multienzyme system from this study, the *B. thuringiensis* spore-based display system would be an excellent model to remediate other persistent organic pollutants in the future.

## RESULTS

### Enzymatic activity of AtzB-bearing spores

The enzymatic activity of AtzB-bearing spores was determined in the reaction with 20 µM (4 mg L^-1^) HA in 25 mM MOPS, pH 7.0 after 12 hours of incubation at room temperature. One mg of *B. thuringiensis* 4Q2-81 spores lacking the *atzB* determinant were included as a control to monitor possible loss of ATR due to adsorption to spore surfaces. The changes in the concentration of HA and NiPA were presented as percentage of that of the initial concentration of HA as illustrated in Fig. 1. More than 35%, 15% and 7% of HA were decreased in the reactions with 1, 0.5, and 0.25 mg AtzB-bearing spores, respectively, in 12 hours (Fig. 1). No production of NiPA was observed in the reactions with *B. thuringiensis* plasmid-free spores (Fig. S1). The mass of AtzB-bearing spores as a function of the degradation of HA was shown in Fig. 1, indicating a positive linear relationship between the enzymatic activity and the mass of AtzB-bearing spores.

**Figure 1.**
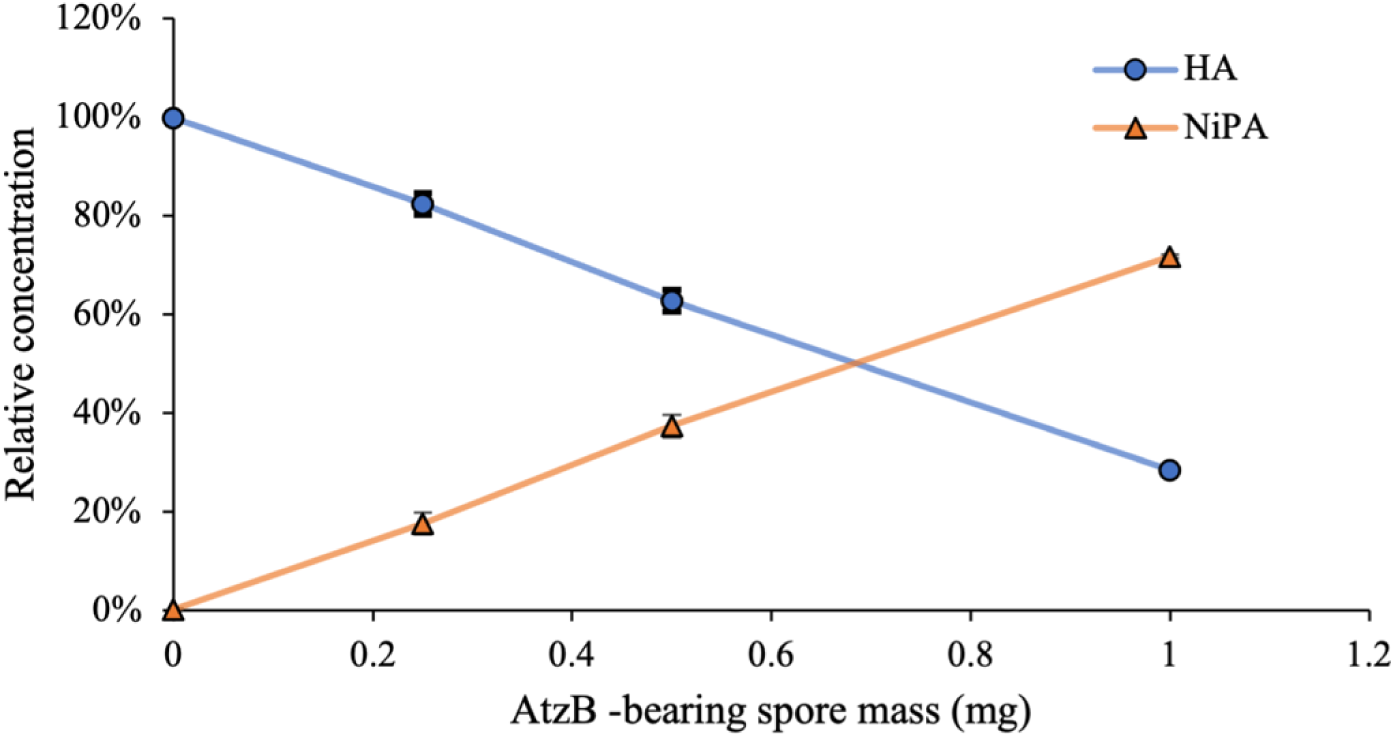
AtzB-bearing spore mass as the function of hydroxyatrazine degradation. Various masses of AtzB-bearing spores were applied to degrade in the reactions with 20 µM (4 mg L^-1^) hydroxyatrazine (HA). Production of HA metabolite, NiPA, was utilized to define the enzymatic activity. Error bars indicate the standard deviation of two independent duplicates. (Blue circle: HA; orange triangle: NiPA)

### Enzymatic activity of AtzB-bearing spores versus recombinant AtzB enzyme

Although the actual number of fused AtzB on the surface of the spore is difficult to determine, the enzymatic activity of AtzB-bearing spores can be determined as the equivalent activity of recombinant AtzB protein. The enzymatic activities of recombinant AtzB protein and AtzB-bearing spores were determined in the reactions with 20 µM (4 mg L^-1^) HA in 25 mM MOPS, pH 7.0 after 12-hour incubation at room temperature. The mass of the recombinant AtzB protein prep as a function of the production of NiPA is shown in Fig.2. The production of NiPA by 0.5 mg AtzB-bearing spores in 12 hours was 3.95 µM. Considering the 31% purity of recombinant AtzB protein prep after ammonium sulfate precipitation (55), 1 mg AtzB-bearing spores exhibited equivalent activity to 514.5 ng pure recombinant AtzB enzyme. Given the molecular weight of AtzB (52,000 g/mol) and 1.2 ×10^8^ spores in 1 mg of a wet pellet determined by plate count, each spore exhibited an activity equivalent to approximately 4.948×10^4^ molecules of AtzB enzyme.

**Figure 2.**
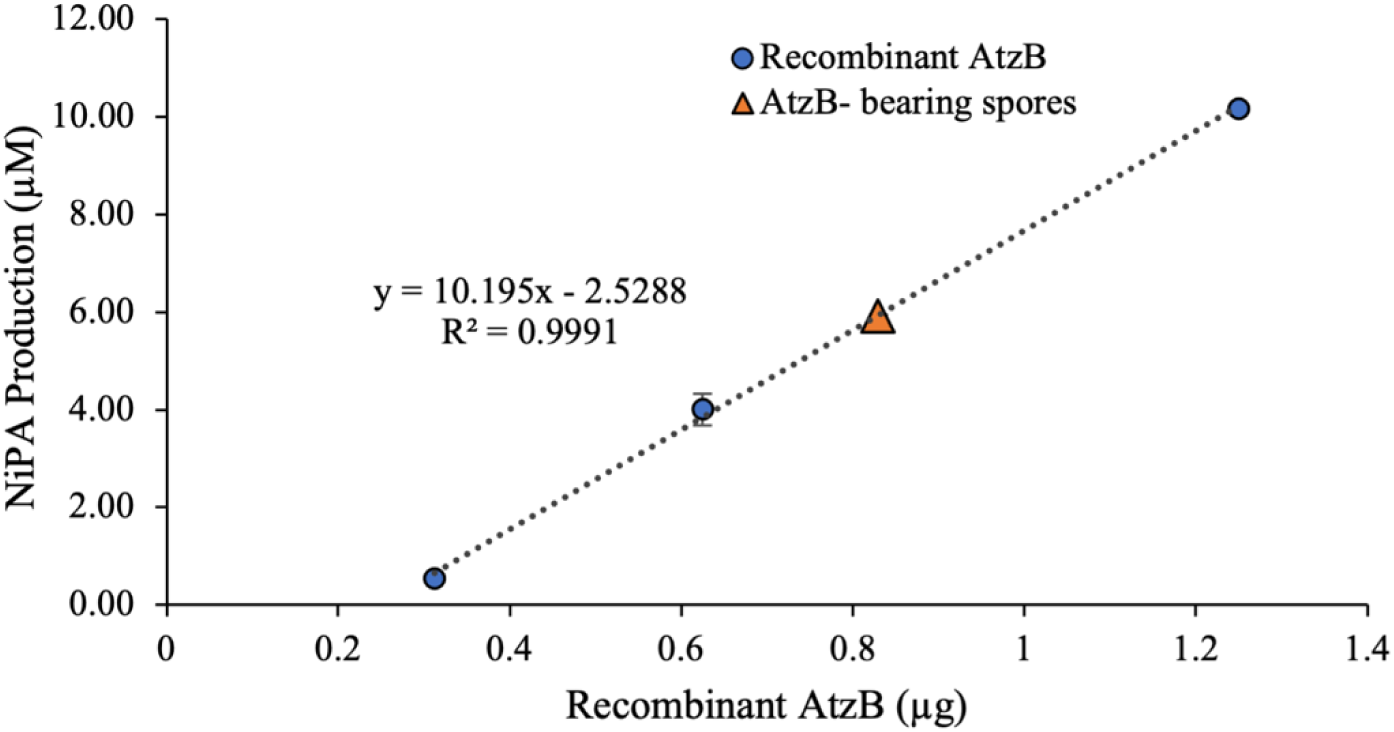
Recombinant AtzB mass as the function of hydroxyatrazine degradation. Various amounts of AtzB recombinant protein in a 1:2 serial dilution or 0.5 mg AtzB-bearing spores were applied in the reactions with 20 µM (4 mg L^-1^) HA. Production of HA metabolite, NiPA, was utilized to define the enzymatic activity. Error bars indicate the standard deviation of two independent duplicates (blue circle: AtzB-bearing spores, orange triangle: Recombinant AtzB protein).

### Enzymatic kinetic assays of AtzB-bearing *B. thuringiensis* spores

The enzymatic kinetic of AtzB spores was determined by the reactions using 0.25 mg AtzB-bearing spores in the reaction with 5, 10, or 20 µM (1, 2, or 4 mg L^-1^) HA in 25 mM MOPS, pH 7.0 in a 12-hour time course study. *B. thuringiensis* plasmid-free spores were used as an indicator for substrate/product absorbance to spores (Fig. S2). It has shown the degradation of HA by AtzB-bearing spores (Fig. 3A) corresponded to the NiPA production (Fig. 3B) over time. The production of NiPA as the function of time demonstrated a linear relationship within the first 8 hours but enzymatic reactions in 5 and 10 µM HA reached the plateau between 8 and 24 hours (Fig. 3B). The enzymatic kinetics K_m_ and V_max_ of 1 mg of AtzB-bearing spores were 25.8 µM and 0.84 nmole NiPA h^-1^ mg^-1^ spores respectively (Table 3) by the calculation with the Michaelis–Menten equation. The AtzB-bearing spores exhibited K_cat_ 1.914 h^-1^ and enzyme efficiency (k_cat_/K_m_) 0.074 µM^-1^h^-1^.

**Figure 3.**
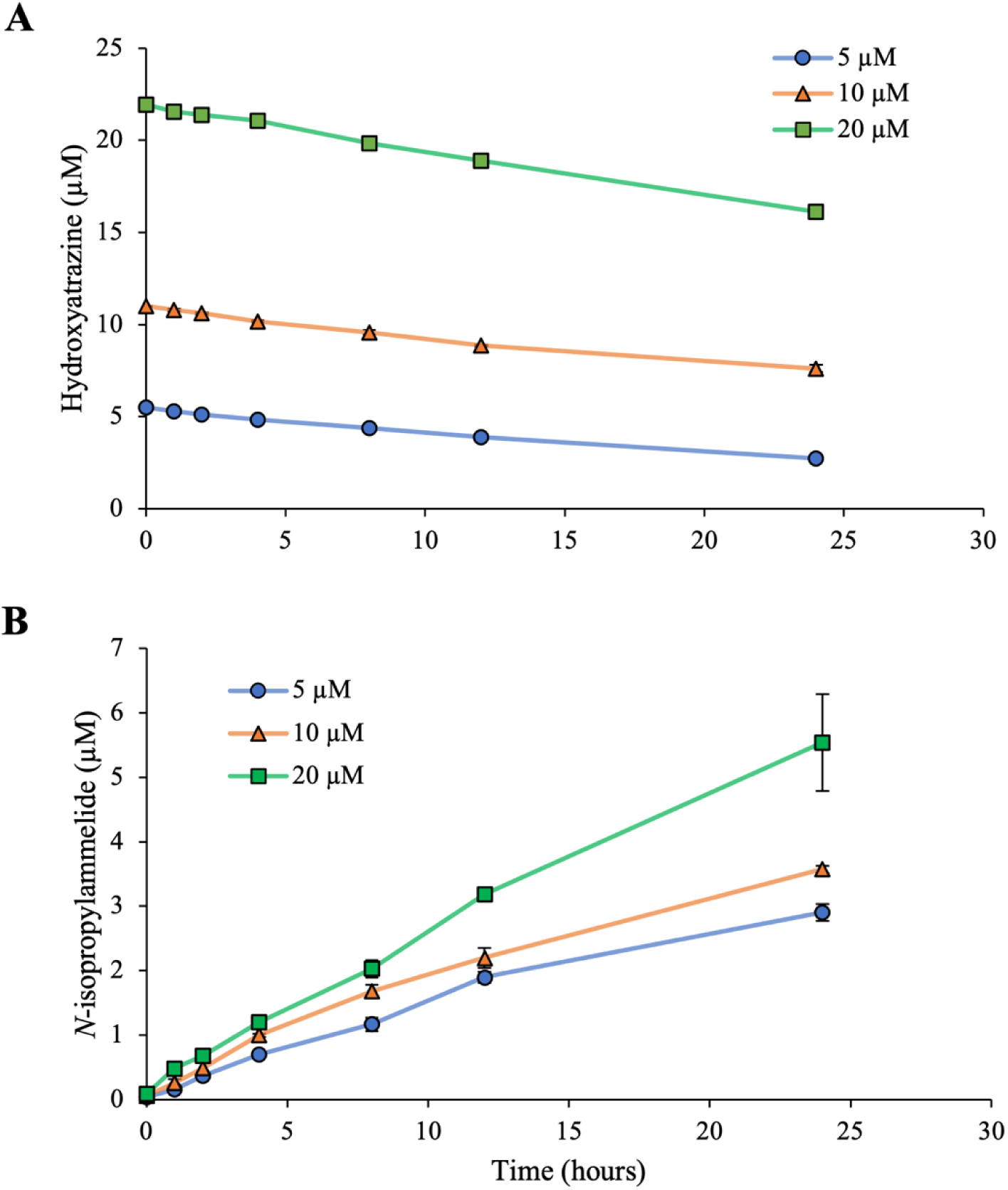
Enzymatic kinetic assay of AtzB-bearing *B. thuringiensis* spores. (A) The degradation of hydroxyatrazine (HA) and (B) production of *N*-isopropylammelide (NiPA). Various concentrations of HA (blue circle: 5 µM, orange triangle: 10 µM, green square: 20 µM) were applied in the reactions with 0.25 mg AtzB-bearing spores in duplicates. Error bars indicate the standard deviation of two independent duplicates.

### Time course study of atrazine degradation in water by AtzA-bearing and AtzB-bearing spores

To establish the optimal ratio of AtzA- to AtzB-bearing spores in one-pot reactions of ATR degradation, a kinetics assay of ATR degradation was performed using 1 mg of AtzA-bearing spores in combination with either 1, 2, or 4 mg of AtzB-bearing spores in the reactions with 46 µM (10 mg L^-1^) ATR in 25 mM MOPS, pH 7.0, in a 96-hour time course study. *B. thuringiensis* plasmid-free spores were used as controls to monitor possible loss of ATR due to adsorption to spore surfaces (Fig. 4C). A background control without spores was used to monitor the possible loss due to evaporation (Fig. 4E). The changes in the concentration of ATR, HA, and NiPA are expressed and presented as percentage of the initial ATR concentration.

**Figure 4.**
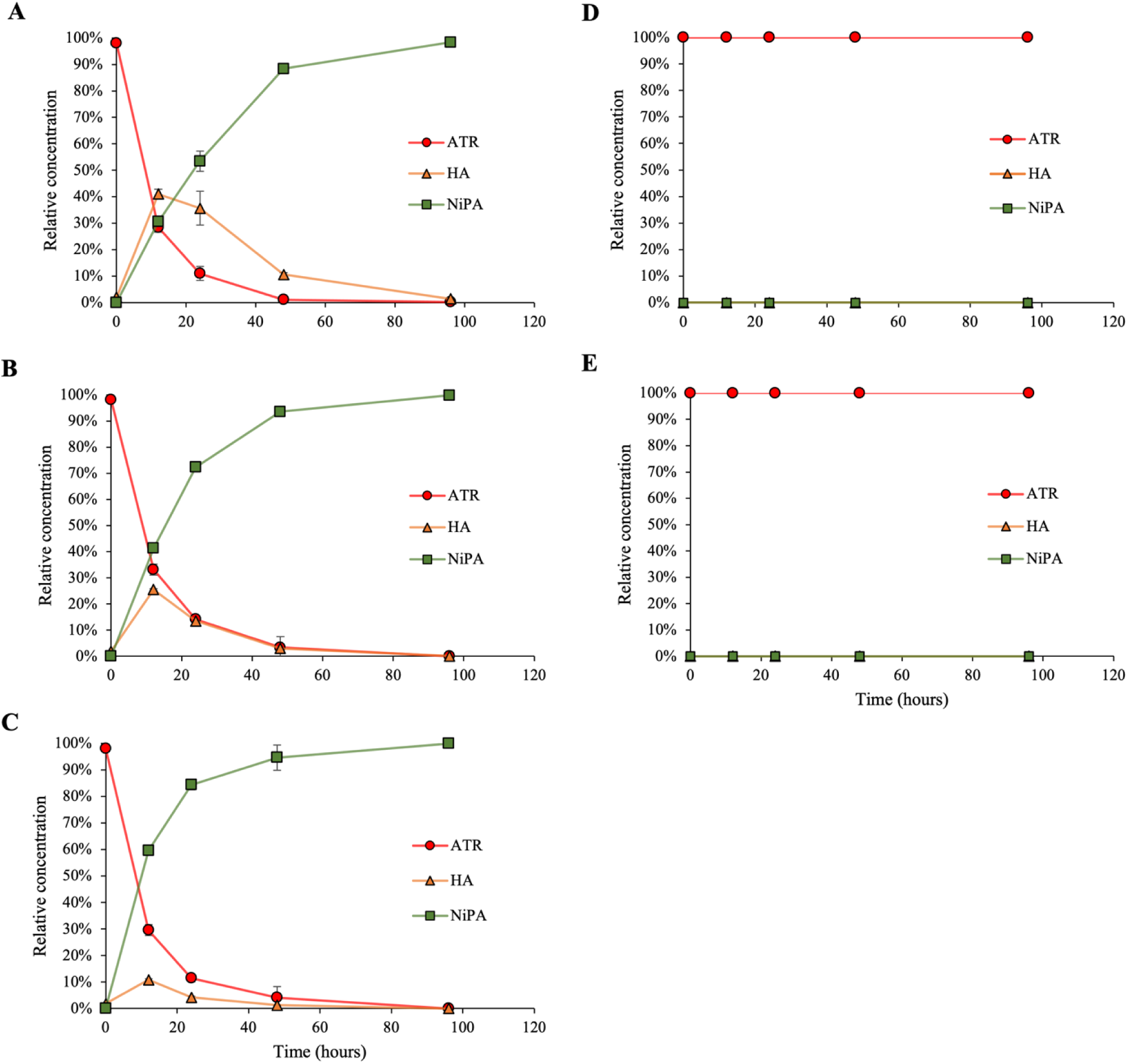
Time as the function of atrazine degradation in water by AtzA-bearing and AtzB-bearing *B. thuringiensis* spores. Spores were pelleted and resuspended in 25 mM MOPS (pH 7.0). (A) 1 mg AtzA-bearing spores and 1 mg AtzB-bearing spores; (B) 1 mg AtzA-bearing spores and 2 mg AtzB-bearing spores; (C) 1 mg AtzA-bearing spores and 4 mg AtzB-bearing spores; (D) 5 mg *B. thuringiensis* plasmid-free spores; and (E) Spore-free control reaction. Atrazine was added to a final concentration of 46 µM (10 mg L^-1^) and MOPS (pH 7.0) to a final concentration of 25 mM. The final concentrations of ATR, HA, and NiPA are expressed as the percentage of the initial ATR, 46 µM (10 mg L^-1^). Error bars indicate the standard deviation of two independent duplicates. (Red circle: ATR, orange triagle: HA, green square: NiPA)

The various ratios of AtzA- to AtzB-bearing spores in one-pot reactions exhibited different rates of HA production/depletion and NiPA production. In the reactions of 1 mg AtzA spores and 1 mg AtzB spores (Fig. 4A), the concentration of HA was generated from 41% of the initial ATR in 12 hours but was reduced rapidly to 10% between 12 and 48 hours, and gradually depleted between 48 and 96 hours. Once HA was generated from ATR degradation, the production of NiPA escalated to almost 90% in 48 hours but gradually reached 100 % between 48 and 96 hours. In the reactions of 1 mg AtzA spores and 2 mg AtzB spores (Fig. 4B), the concentration of HA was shown as 25 % of the applied ATR in 12 hours, decreased to 5% between 12 and 48 hours, and reached complete depletion between 48 and 96 hours. The production of NiPA escalated to above 70% in 24 hours but reached 100% between 48 and 96 hours. In the reactions of 1 mg AtzA spores and 4 mg AtzB spores (Fig. 4C), the concentration of HA was shown as 10% in 12 hours and gradually degraded between 12 and 48 hours. The production of NiPA escalated to 85% in 24 hours but gradually reached 100% between 48 and 96 hours. The applied ATR in all the reactions with AtzA and AtzB-bearing spores was completely transformed to NiPA in 96 hours. No ATR loss by absorption or evaporation was observed in the control reactions (Fig. 4D and 4E).

### Time course study of atrazine degradation in surface water by AtzA- and AtzB-bearing spores

To simulate the remediation in surface water, ATR degradation assays with 1 mg of AtzA-bearing spores and 2 mg of AtzB-bearing spores in surface water, collected from the Missouri River, fortified with ATR for the final concentration of 34.5 nM (7.5 µg L ^-1^) were performed in a 96-hour time course study. *B. thuringiensis* plasmid-free spores were used as controls to monitor possible loss of ATR due to adsorption to spore surfaces. A background control without spore was used to determine the existing ATR in surface water and the possible loss of ATR due to evaporation. Relatively final concentrations of ATR, HA, and NiPA are presented as a percentage of the initial ATR concentration. As illustrated in Fig. 5A, approximately 90% of applied ATR was converted to 20% HA and 70% NiPA in 24 hours. More than 80% of NiPA production was observed in the reactions in 48 hours. The applied ATR was completely metabolized to HA and further converted to NiPA in the reactions in surface water in 96 hours. No ATR loss by absorption was observed in the spore control reactions (Fig 5B). Existing ATR and HA were observed in the background control reactions from the start, but no depletion of ATR and HA happened due to the evaporation over time (Fig. 5C).

**Figure 5.**
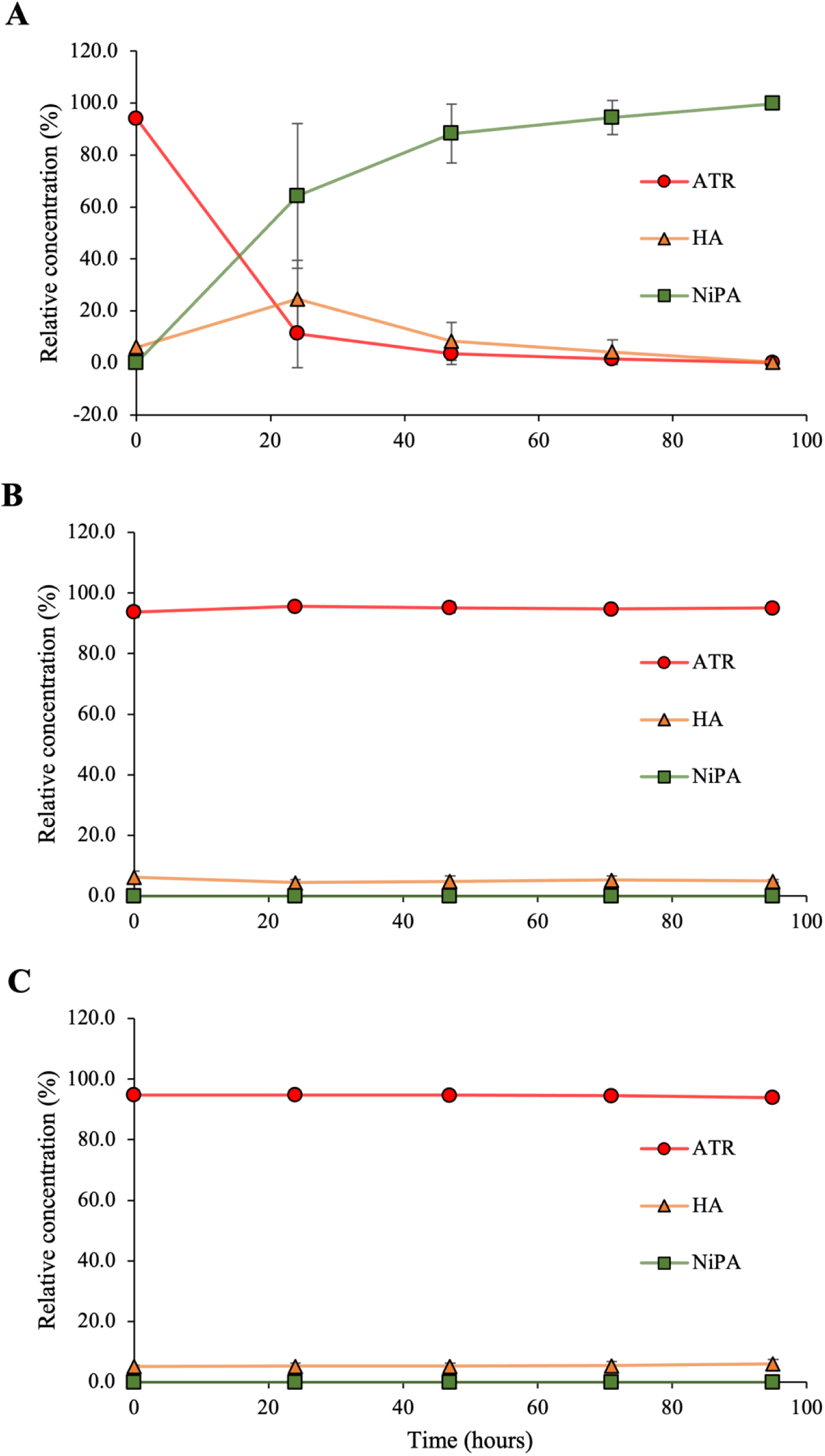
Time as the function of atrazine degradation in surface water. Spores were resuspended in lake water fortified with ATR at a final concentration of 34.5 nM (7.5 µg L ^-1^). (A) 1 mg AtzA-bearing spores and 2 mg AtzB-bearing spores; (B) 3 mg *B. thuringiensis* plasmid-free spores; (C) no spores. The final concentration of ATR, HA, and NiPA are expressed as the percentage of the applied ATR, 34.5 nM (7.5 µg L^-1^). Error bars represented the standard deviation of two independent experiments. (Red circle: ATR, orange triangle: HA, green square: NiPA)

### Chemical characterization of surface water

The chemical characterization of the Missouri River surface water utilized in this study showed 43.5 mg L^-1^ calcium (Ca^2+^), 23.8 mg L^-1^ magnesium (II), 0.001 mg L^-1^ zinc (II), 0.003 mg L^-1^ iron (II), 0.013 mg L^-1^ manganese (II), 0.016 mg L^-1^ copper (II) as well as 0.007 mg L^-1^ aluminum (III) and a pH value 8.22 (Table 4).

## DISCUSSION

In this study, the *B. thuringiensis* spore-based display system successfully expressed the second enzyme in the atrazine degradation pathway, AtzB. Hydroxyatrazine *N*-ethylaminohydrolase (AtzB), the only known enzyme that transforms HA to NiPA, plays a critical role in ATR degradation (55). NiPA can be further transformed into cyanuric acid, the central intermediate in ATR metabolism, that can be metabolized by many other bacteria (55). Also, the ratio of applied AtzA- and AtzB-bearing spores was optimized to achieve 100% conversion of ATR, 46 µM (10 mg L^-1^) to NiPA in a one-pot multienzyme reaction as well as 100% transformation of ATR, 34.5 nM (7.5 µg L^-1^) in the surface water to NiPA in 96 hours. These findings validate that the *B. thuringiensis* spore-based display system could be utilized for as the platform to catalyze the multi-step enzymatic reactions. The spore nature of this system bypasses bioremediation challenges, such as the requirement for nutrient supplementation to permit bacterial growth.

Characterization of AtzB tethered to the surface of *B. thuringiensis* spores is important before the introduction of AtzB-bearing spores into multienzyme reactions. The enzymatic activity, deamination of HA to NiPA (54), of AtzB tethered on the spores was not lost due to the addition of BclA targeting domain (28, 34) at the N terminus or the anchoring of the fusion protein to the spore surface (Fig. 1). It suggested that the configuration of enzymatically active site of AtzB was not altered by the N-terminal fusion of BclA. The kinetic values, K_m_ and K_cat_, of AtzB-bearing spores were determined as 25.8 µM and 26.31 h^-1^, respectively (Table 3) in comparison to the K_m_ and K_cat_ of free AtzB protein were 20 µM and 3.2 s^-1^ (55). Due to its higher K_m_ values, AtzB-bearing spores have a slightly weaker affinity for substrate than free AtzB enzyme. Besides, the K_cat_ of AtzB-bearing spores was much lower than free AtzB protein, as immobilization platforms and techniques often greatly impact the K_cat_ of immobilized enzymes (44, 56, 57). In the *B. thuringiensis* spore display system, the other native surface proteins on the spores might cause less accessibility of fused AtzB enzyme to its substrate, which could result in lower K_cat_ of AtzB-bearing spores than free AtzB enzyme. However, because the production of AtzB-bearing spores is a simpler, less time-consuming procedure, and substantially less costly process than that of purifying AtzB enzyme, high K_m_ and low K_cat_ of AtzB-bearing spores could be easily compensated by applying a proper mass of spores to achieve similar HA degradation to purifying AtzB enzyme.

Each AtzA-bearing spore was previously estimated to exhibit the activity equivalent to 2.076×10^3^ AtzA enzyme molecules (34) and each AtzB-bearing spore was estimated in this study to exhibit the activity equivalent to 4.948×10^4^ AtzB enzyme molecules. The difference could result from the K_m_ and K_cat_ values of AtzA- and AtzB-bearing spores. (Table 3). According to K_m_ and K_cat_ values in enzymatic kinetics, a greater mass of AtzB-bearing spores would be needed in the multienzyme reaction in a one-pot reaction. Furthermore, AtzA-bearing spores exhibited slightly higher turnover (K_cat_) than AtzB-bearing spores (Table 3), suggesting that once ATR was converted to HA by AtzA-bearing spores, there might be a delayed initiation of the reaction from AtzB-bearing spores until enough HA accumulated to pass the threshold in degradation of HA to NiPA. Finally, the value of K_cat_/K_m_ indicates how effectively the enzyme captures its substrate for catalytic turnover (58). AtzB-bearing spores exhibited higher K_cat_/K_m_, approximately 7-fold to that of AtzA-bearing spores, suggesting AtzB-bearing spores could have higher enzymatic degradation rates than AtzA-bearing spores at low concentrations of substrates.

The applied ATR, 46 µM (10 mg L^-1^), was completely converted to the end metabolite, NiPA, in one-pot reactions using two enzymes delivered by *B. thuringiensis* spore display system in surface water at benchtop with various ratios (1:1, 1:2, 1:4) of AtzA- to AtzB-bearing spores (Fig.4 A-C). The HA production peaked in 12 hours in all reactions, but the depletion of HA was only delayed in the reactions with 1 mg of AtzB-bearing spores until 96 hours (Fig. 4A). Although the HA depletion and NiPA production in the reactions with 4 mg AtzB-bearing spores initiated sooner in 24 hours (Fig. 4C) than that in the reactions with 2 mg AtzB-bearing spores in 48 hours (Fig. 4B), more than 90% of applied ATR was converted to NiPA in both treatments in 48 hours. Therefore, the 1:2 ratio of applied AtzA- to AtzB-bearing spores in the reaction would be sufficient to achieve the complete degradation of ATR to NiPA.

The highest ATR concentration of 34.5 nM (7.5 µg L^-1^) in the surface water, observed in Nebraska, U.S.A. in the past two decades (59), was simulated by fortifying ATR in surface water in this study. This could be the first benchtop study using multienzymes delivered by the *B. thuringiensis* spore display system with 34.5 nM (7.5 µg L^-1^) ATR in the surface water. The results showed approximately 90% of the applied ATR, 27.6 nM (6 µg L^-1^), was degraded in 72 hours. Most importantly, over 80% of applied ATR was degraded and the concentration of ATR dropped below U.S. EPA MCL (3 µg L^-1^ in drinking water) in 24 hours. The applied ATR was completely converted to NiPA in 96 hours (Fig. 5A). Therefore, the *B. thuringiensis* spore display system demonstrated the promising potential for degrading ATR in drinking water sources.

The *B. thuringiensis* spore display system overcame the major challenges of enzyme-based bioremediation in the environment, including stability of enzymatic activity, and no requirement for cofactor supplementation (60). Previously, an enzyme-based remediant developed by CSIRO Ecosystem Science showed a half-life of 79 hours in the natural water (61). Herein, both AtzA- and AtzB-bearing spores exhibited enzymatic activity in the surface water collected from the Missouri River for up to 96 hours, indicating a longer stability of enzymatic activity in natural water. Additionally, AtzB is a dimeric enzyme from the amidohydrolase superfamily and is reported to contain 1 zinc molecule per monomer, which is essential for catalytic activity (55). However, because the production of AtzB-bearing spores is a simpler, less time-consuming procedure, and substantially less costly process than that of purifying free AtzB enzyme, the bioremediation process using this platform remains economically feasible.

Furthermore, enzymatic activities of AtzA and AtzB expressed on the *B. thuringiensis* spore display system were not affected by other nitrogen sources present in environments. Results from Bichat *et al.* showed that *Pseudomonas* sp. strain ADP previous growth on nitrate (NO_3_^-^), ammonium(NH_4_^+^), or urea exhibited reducing rate of ATR degradation in soil (62) García-González *et al.* also reported that nitrate amendment (0.5 mg of NO_3_N g^-1^) in soil resulted in decreased atrazine mineralization by *Pseudomona*s sp. strain ADP(46). However, our results showed, regardless of the nitrate level (1. 71 mg L^-1^) in the surface water collected from Missouri River, both AtzA- and AtzB-bearing spores consistently degraded ATR to NiPA throughout 96-hour incubation (Fig. 5A), suggesting no down-regulation of ATR degradation enzymes. Therefore, because the spore platform functions without requiring growth of the *B. thuringiensis*, there is no induction of gene expression required and thus no inhibition resulting from the presence of nitrogen sources in the environment.

In summary, our findings suggest that the *B. thuringiensis* spore display system has demonstrated great potential as a bioremediation technology for degrading ATR in drinking water sources for the following reasons: (1) the flexibilities in enzyme display and the application of multiple enzymes, (2) no requirement for cofactor supplementation in the enzymatic reaction, (3) stability of enzymatic activities in natural water, and (4) no down-regulation of ATR enzymes was observed. For future studies, the next enzyme in the ATR degradation, AtzC, will be investigated and incorporated into the multienzymes reaction using the *B. thuringiensis* spore display system. These three enzymes delivered by the spore display system in one-pot reactions will be an excellent model for the bioremediation of organic persistent pollutants which often require more than one enzyme to complete the detoxification/metabolization process.

## MATERIALS AND METHODS

The brain heart infusion (BHI) broth (Bacto™), Nutrient broth (Oxoid), Luria-Bertani (LB) broth, 0.2 µm PES membrane filter (Thermo Scientific, cat# 09-741-07), HPLC-grade methanol and acetonitrile were purchased from Fisher Scientific, U.S.A. The 0.45 µm analytical test filter funnel (#145-0045) for harvesting competent cells and the Whatman 0.2 µm Anotop syringe membrane filter were purchased from Thermo Fisher, U.S.A. The Wizard™ Plus SV Minipreps DNA Purification System and Wizard™ Genomic DNA Purification Kits were purchased from Promega, U.S.A. The StrataClone PCR cloning kits were purchased from Agilent Technologies, U.S.A. The 0.2 cm gap cuvettes for electroporation were purchased from Midwest Scientific, U.S.A. The analytical standards of atrazine (≥98%) and hydroxyatrazine (≥98%) were purchased from Millipore Sigma, St. Louis, U.S.A., and *N*-isopropylammelide was kindly gifted from Dr. Lawrence P. Wackett at the University of Minnesota. The Kinetex C18 (100mm x 4.6 mm; 2.6 µm particle size) reverse-phase column was purchased from Phenomenex, U.S.A.

### Bacterial strains and growth conditions

*Pseudomonas sp*. strain ADP was grown in R-medium containing 0.1% (wt/vol) sodium citrate as the carbon source and atrazine as the only nitrogen source at a final concentration of 0.46 mM (100 µg L^-1^), at 30°C with agitation (36, 37). *Escherichia coli* (DH5α, GM48, and SoloPack®) were grown in Luria-Bertani (LB) broth containing appropriate antibiotics for selection at 37°C with agitation for the procedure of cloning or protein production. A crystal-negative strain of *Bacillus thuringiensis* subsp. *israelensis* (*Bacillus* Genetic Stock Center 4Q2-81) (63) was grown in BHI broth for competent cell preparation in electroporation. AtzA-bearing spore culture was grown in LB broth + 10 mg mL^-1^ Chloramphenicol at 30°C and AtzB-bearing spore culture was grown in Nutrient broth containing 10 mg mL^-1^ Chloramphenicol at 25°C both with 225 rpm agitation.

### DNA Extraction

Overnight cultures of plasmid-containing *E. coli, B. thuringiensis* subsp. *israelensis* (4Q2-81) and *Pseudomonas sp*. strain ADP were harvested by centrifugation. The plasmid DNA in *E. coli* was isolated using Wizard™ Plus SV Minipreps DNA Purification System, following the standard protocol provided by Promega. The total DNA of *B. thuringiensis* subsp. *israelensis* (4Q2-81) and *Pseudomonas sp*. strain ADP were extracted using Wizard™ Genomic DNA Purification Kits following the standard protocol provided by Promega.

### Cloning of the *atzB* determinants to the spore expression vector

Plasmid pADP-1 DNA was isolated from an overnight culture of *Pseudomonas sp.* strain ADP. The *atzB* determinant was PCR amplified using primers 5’-ctcgagATGACCACCACTCTTTACACC-3’ and 5’--ctcgagTCAGCATGGTGTGACACCGG-3’, which incorporated *XhoI* site at 5’ end (shown in lowercase letters). The PCR cycling conditions were 94°C for 3 min and 35 cycles of 94°C for 30 s, 62 °C for 30 s, 58°C for 30 s, and 72°C for 90 s. The 1.44-kb PCR fragment was cloned into the StrataClone PCR cloning vector pSC-A-amp/kan plasmid and the nucleotide sequence of the cloned fragment was verified at the University of Missouri DNA Core Facility. The *atzB* open reading frame was further cloned as an *Xho*I fragment into plasmid pHH4283 (34) (Table 1), the *B. thuringiensis* spore expression vector (34). The correct sequence and orientation of the insert were verified by DNA sequencing at the University of Missouri DNA Core and the plasmid was designated as pHH4295.

**Table 1.**
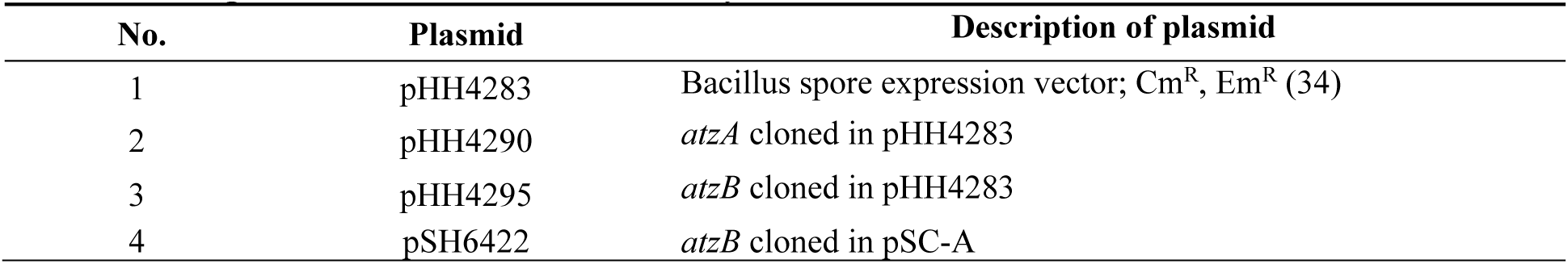
The plasmids described in this study.

| No. | Plasmid | Description of plasmid |
| --- | --- | --- |
| 1 | pHH4283 | Bacillus spore expression vector; Cm<sup>R</sup>, Em<sup>R</sup> (34) |
| 2 | pHH4290 | *atzA* cloned in pHH4283 |
| 3 | pHH4295 | *atzB* cloned in pHH4283 |
| 4 | pSH6422 | *atzB* cloned in pSC-A |

### Electroporation of *B. thuringiensis*

The *B. thuringiensis* subsp. *israelensis* (4Q2-81) was grown in BHI+0.5% glycerol broth at 37°C with agitation until the OD_600_ reached 0.6. The competent cells were harvested by filtration through a 0.45 µm analytical test filter funnel, followed by washing with ice-cold electroporation buffer (1 mM HEPES, 10% glycerol, pH 7.0; filter sterile) twice, and resuspended in ice-cold electroporation buffer. One µg of DNA in 10 µl nuclease-free dH_2_O and 100 µl of cell resuspension were transferred into 0.2 cm gap cuvettes. The electroshock was applied using a BioRad Gene Pulser apparatus at 2.25 kV, 25 μF, and 100 Ohms, followed by transfer to a 15 ml polypropylene conical tube and the addition of 1 ml of BGGM broth (BHI with 10% glycerol, 0.4% glucose, and 10 mM MgCl_2_) to revive the shocked cells. The post-electroshocked cells were incubated at 30°C with agitation for 2 hours before plated on BHI plates containing 10 mg mL^-1^ chloramphenicol. The plates were incubated at 30°C overnight.

### Bacillus thuringiensis spore preparation

AtzA-bearing spores were generated by culturing *B. thuringiensis* carrying pHHY4290 (34) (Table 1) in LB with chloramphenicol at 10 mg L^-1^ and AtzB-bearing spores were generated from the *B. thuringiensis* carrying pHH4295 (Table 1) in Nutrient broth with chloramphenicol at 10 mg L^-1^. After 4-7 days of incubation at 25°C, the spores (>90%) were harvested by centrifugation for 1 minute, 2,000 x *g*, and cell debris was removed by washing with phosphate-buffered saline (PBS) repeated 3 to 4 times. The AtzA- and AtzB-bearing spores were normalized to 100 mg mL^-1^ in 25 mM 3-(N-morpholino)propanesulfonic acid (MOPS) buffer, pH 6.9, or PBS, respectively, and stored at 4°C until analyzed for enzymatic activity.

### AtzB protein production and purification

The 1. 97 kb DNA fragment, including the *atzB* promoter and its ORF (54), was amplified by PCR using pADP-1 as a template with primers 5’-AGCCTTGAT CATGAAGGCG AGCATGGTG-3’ and 5’-TCAGCATGGTGTGACACCGGTGCCCCGTC-3’. The PCR-produced fragments were cloned into a PCR cloning vector pSC-A-amp/kan plasmid, creating plasmid pHS6422 (Table 1). The *E. coli* culture carrying pHS6422 was inoculated in 1 L of LB broth with 100 mg L^-1^ ampicillin and continuously grown at 37°C for four hours after OD_600_ reached 0.8. The cells were harvested by centrifugation for 10 min, 3,000 x *g*, and the pellets were resuspended in 25 mM MOPS buffer, pH 7.0 after being frozen at -80°C for 2 hours. The cells were lysed by bead beating with 0.1-mm glass beads for 3 rounds of 1 min at 4°C with 1-min cooling interval, followed by centrifugation for 5 mins at 15,000 x *g* for 5 mins to collect the supernatant as crude extract. The crude extract was collected and solid (NH_4_)_2_SO_4_ was added to achieve 30% saturation. The precipitation was incubated for at least 4 hours or until the salt completely dissolved at 4°C with stirring and then centrifuged for 5 mins at 15,000 x *g* at 4°C to collect protein pellets. The pellets were resuspended in 25 mM MOPS, pH 7.0, and dialyzed against 25 mM MOPS, pH 7.0 to remove the salt at 4°C. The protein preparation was retrieved after dialysis.

### Enzymatic activity assay with AtzB-bearing spores and recombinant AtzB

Twenty µM (4 mg L^-1^) of hydroxyatrazine (HA) in 25 mM MOPS, pH 7.0 was prepared as stock solution. One mL of HA stock solution was added into 2 ml tubes containing 1 mg of AtzB-bearing spores or 10 mL of recombinant AtzB protein suspension in a (1:2) serial dilution in 25 mM MOPS buffer, pH 7.0, in duplicates. The reaction mixtures were incubated at 25°C for 12 hours on a rocking platform. At the end of 12-hour reaction, 490 µl of methanol was added to stop the reactions and to extract HA and its metabolite, NiPA. The extracted mixtures were centrifuged at 15,000 x *g* for 1 min to remove spores, followed by filtering through the Whatman 0.2 µm Anotop syringe membrane filter into a high-performance liquid chromatography (HPLC) injection vial. The vials were stored at -20°C until quantification of HA and NiPA using HPLC-triple quadrupole mass spectrometry (HPLC-MS/MS).

### Enzymatic kinetic study of AtzB-bearing *B. thuringiensis* spores

One ml of 5 µM, 10 µM or 20 µM of HA in 25 mM MOPS, pH 7.0 was applied to two 2-ml tubes containing 250 µg of AtzB-bearing spores as sample reactions and to two 2-ml tubes containing 250 µg of *B. thuringiensis* plasmid-free spores as reaction controls, or 25 mM MOPS (pH 7.0) only as background controls. The reaction mixtures were incubated at 25°C in the dark for a time course of 0, 1, 2, 4, 8, 12, and 24 hours with rocking. At each time point, 490 µL methanol was added to each tube to stop the reaction and to extract HA and the metabolite, NiPA. The extracted mixtures were centrifuged at 15,000 x *g* for 1 min to remove spores, followed by filtering through the Whatman 0.2-m Anotop syringe membrane filter into HPLC injection vials. The vials were stored at -20°C until chemical analysis with HPLC-MS/MS. The final concentration of NiPA was used to determine the K_m_ and V_max_ of AtzB spores using the Michaelis–Menten equation.

### Atrazine degradation by a combination of AtzA-bearing and AtzB-bearing spores

Twenty-three µM (5 mg L^-1^) of atrazine (ATR) in 25 mM MOPS, pH 7.0, was prepared as ATR stock solution. One mg, 2 mg, or 4 mg of AtzB-bearing spores was added to two 2-ml tubes containing 1 mg of Atz-bearing spores, followed by adding 1 ml of ATR stock solution to set up the reactions. One ml of ATR stock solution was applied to two 2 ml tubes containing 5 mg of *B. thuringiensis* plasmid-free spores as reaction control or 25 mM MOPS, pH 7.0 only as background control. The reaction mixtures were incubated at 25°C for a time course of 0, 12, 24, 48, and 96 hours with gentle rocking. At each time point, 500 µL methanol was added to stop the reaction and to extract ATR, HA, and NiPA. The extracted mixtures were centrifuged at 15,000 x *g* for 1 min to remove spores, followed by filtering through the Whatman 0.2 µm Anotop syringe membrane filter into HPLC injection vials. The vials were stored at -20°C until quantification of ATR, HA, and NiPA using HPLC-MS/MS.

### Atrazine degradation in surface water by a combination of AtzA-bearing and AtzB-bearing spores

The surface water collected from the Missouri River was sterilized by passing it through a 0.2 µm filter. The sterilized surface water was fortified with ATR to achieve 34.5 nM (7.5 µg L^-1^) as the final concentration, which was the highest concentration of ATR detected in surface water in the past two decades (59). One mg of AtzA-bearing spores and 2 mg AtzB-bearing spores were added in each of the 2 ml tubes containing 1 ml of ATR-fortified surface water to set up the reactions. 1 ml of ATR-fortified surface water was added to 2 brown bottles containing 3 mg of *B. thuringiensis* plasmid-free spores as reaction control or 25 mM MOPS, pH 7.0 only as background control. The reaction mixtures were incubated at 25°C for a time course of 0, 24, 48, 72, and 96 hours with rocking. At each time point, 500 µL methanol was added to stop the reaction and to extract ATR, HA, and NiPA. The extracted mixtures were centrifuged at 15,000 x *g* for 1 min to remove spores, followed by filtering through the Whatman 0.2 µm Anotop syringe membrane filter into HPLC injection vials. The vials were stored at -20°C until quantification of ATR, HA, and NiPA using HPLC-MS/MS.

### Chemical analyses. (NEED Dr. Lin to review)

The quantification of ATR, HA, and NiPA was performed using a Waters Alliance 2695 High-Performance Liquid Chromatography (HPLC) system coupled with a Waters Acquity TQ triple quadrupole mass spectrometer (MS/MS). The analytes were separated using a Kinetex C18 (100mm x 4.6 mm; 2.6 µm particle size) reverse-phase column. The mobile phase consisted of (A) 0.1% formic acid in water and (B) 100% acetonitrile. The gradient conditions were 0 – 0.3 min, 2% B; 0.3-7.27 min, 2-80% B; 7.27-7.37 min, 80-98% B; 7.37-9.0 min, 98% B; 9-10 min 98-2% B; 10.0 – 15.0 min, 2% B at the flow rate of 0.5 mL/min. The ion source in the MS/MS system was electrospray ionization (EI) operated in either positive or negative ion mode with a capillary voltage of 1.5 kV. The ionization sources were programmed at 150°C and the desolvation temperature was programmed at 450°C. The optimized collision energy, cone voltage, molecular and product ions of ATR and its metabolites are summarized in Table 2.

**Table 2.**
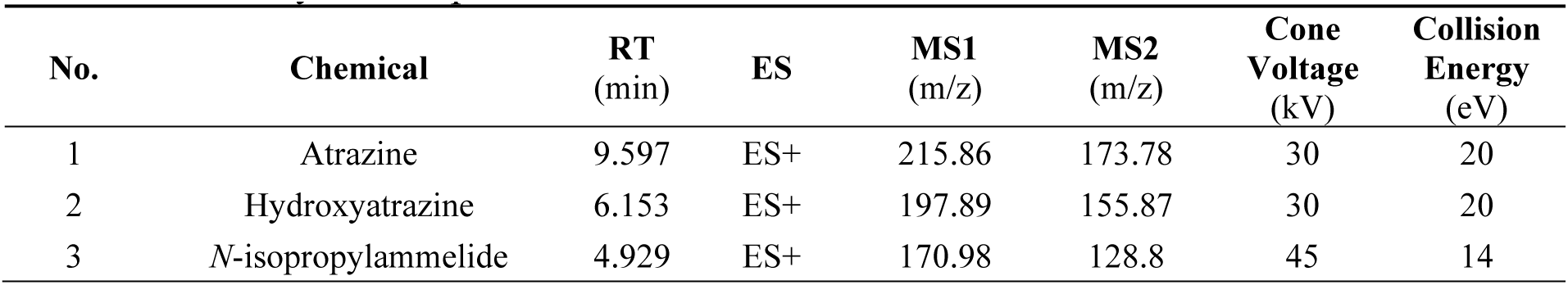
Summary of the optimized LC-MSMS Parameters for ATR and its metabolites.

| No. | Chemical | RT<br>(min) | ES | MS1<br>(m/z) | MS2<br>(m/z) | Cone<br>Voltage<br>(kV) | Collision<br>Energy<br>(eV) |
| --- | --- | --- | --- | --- | --- | --- | --- |
| 1 | Atrazine | 9.597 | ES+ | 215.86 | 173.78 | 30 | 20 |
| 2 | Hydroxyatrazine | 6.153 | ES+ | 197.89 | 155.87 | 30 | 20 |
| 3 | *N*-isopropylammelide | 4.929 | ES+ | 170.98 | 128.8 | 45 | 14 |

**Table 3.**
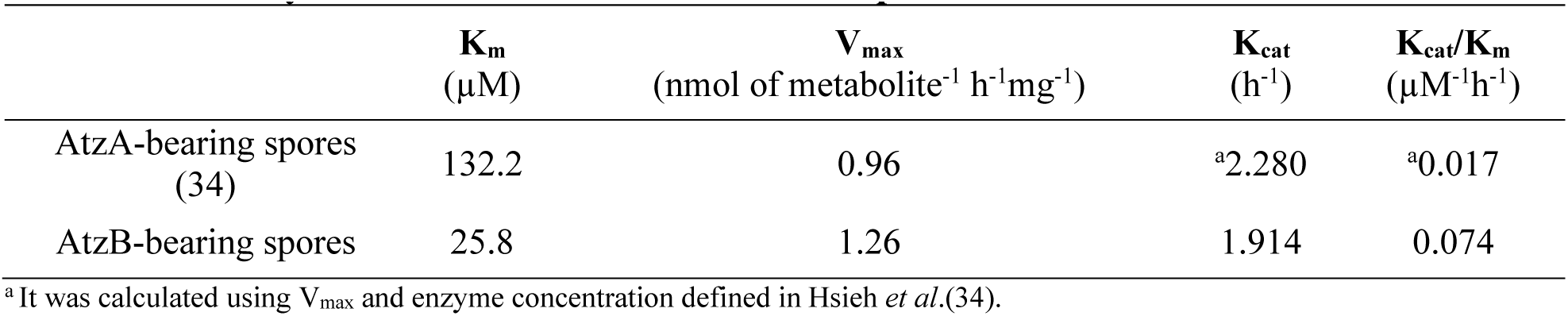
The enzymatic kinetics of AtzA and AtzB spores.

| | **K<sub>m</sub>**<br>($\mu\text{M}$) | **V<sub>max</sub>**<br>(nmol of metabolite<sup>-1</sup> h<sup>-1</sup>mg<sup>-1</sup>) | **K<sub>cat</sub>**<br>(h<sup>-1</sup>) | **K<sub>cat</sub>/K<sub>m</sub>**<br>($\mu\text{M}^{-1}\text{h}^{-1}$) |
| --- | --- | --- | --- | --- |
| AtzA-bearing spores<br>(34) | 132.2 | 0.96 | <sup>a</sup>2.280 | <sup>a</sup>0.017 |
| AtzB-bearing spores | 25.8 | 1.26 | 1.914 | 0.074 |

<sup>a</sup> It was calculated using V<sub>max</sub> and enzyme concentration defined in Hsieh *et al.*(34).

**Table 4.** The water analysis of surface water collected from the Missouri River.

| Tests | Units | Results |
| --- | --- | --- |
| pH |  | 8.22 |
| Electrical Conductivity (E. C.) | mmho/cm | 0.621 |
| Nitrate – N (NO<sub>3</sub> - N) | mg/l | 1.71 |
| Ammonium-N (NH<sub>4</sub>-N) | mg/l | 0.218 |
| Phosphate (PO<sub>4</sub><sup>3-</sup>) | mg/l | 0.145 |
| Potassium (K<sup>+</sup>) | mg/l | 8.54 |
| Calcium (Ca<sup>2+</sup>) | mg/l | 43.5 |
| Magnesium (Mg<sup>2+</sup>) | mg/l | 23.8 |
| Sodium (Na<sup>+</sup>) | mg/l | 52.3 |
| Zinc (Zn<sup>2+</sup>) | mg/l | 0.001 |
| Iron (Fe<sup>2+</sup>) | mg/l | 0.003 |
| Manganese (Mn<sup>2+</sup>) | mg/l | 0.013 |
| Copper (Cu<sup>2+</sup>) | mg/l | 0.016 |
| Chloride (Cl<sup>-</sup>) | mg/l | 6.46 |
| Sulfate (SO<sub>4</sub><sup>2-</sup>) | mg/l | 175 |
| Bicarbonate (HCO<sub>3</sub><sup>-</sup>) | mg/l | 88 |
| Carbonate (CO<sub>3</sub><sup>-</sup>) | mg/l | 5 |
| Total Dissolved Solids | mg/l | 397 |
| Hardness mg/l | mg/l | 207 |
| SAR |  | 1.58 |
| Boron mg/l | mg/l | 0.071 |
| Aluminum (Al<sup>+++</sup>) | mg/l | 0.007 |
| Silica (SiO<sub>2</sub>) mg/l | mg/l | 0.003 |
| Alkalinity | mg/l | 94 |

### Surface water collection and characterization

The surface water of the Missouri River was collected from the Boonville drinking water treatment plant on July 9^th^, 2019, shipped at 4°C and stored at -20°C. The sample was filtered through a 0.2 µm PES membrane filter before they were characterized at the University of Missouri Soil and Plant Testing Lab (https://extension.missouri.edu/programs/soil-and-plant-testing-laboratory/spl-water-analysis).

## ACKNOWLEDGMENTS

This research was funded by a grant from Mizzou Advantage and support from the University of Missouri Center for Agroforestry and USDA/ARS Dale Bumpers Small Farm Research Center under agreement number 58-6020-6-001 from the USDA Agricultural Research Service. Thank Dr. Lawrence P. Wackett at the University of Minnesota for kindly gifting the *N*-isopropylammelide as a chemical standard.

